# Proximity to human settlement is directly related to carriage of critically important antimicrobial-resistant *Escherichia coli* and *Klebsiella pneumoniae* in Silver Gulls

**DOI:** 10.1101/2021.09.17.460878

**Authors:** Shewli Mukerji, Shafi Sahibzada, Rebecca Abraham, Marc Stegger, David Jordan, David J Hampson, Mark O’Dea, Terence Lee, Sam Abraham

## Abstract

Human population and activities play an important role in dissemination of antimicrobial resistant bacteria. This study investigated the relationship between carriage rates of critically important antimicrobial-resistant (CIA-R) *Escherichia coli* and *Klebsiella pneumoniae* by Silver Gulls and their proximity to human populations. Faecal swabs (n=229) were collected from Silver Gulls across 10 southern coastline locations in Western Australia (WA). The sampling locations included main town centres and remote areas. Fluoroquinolone and extended-spectrum cephalosporin-resistant *E. coli* and *K. pneumoniae* were isolated and tested for antimicrobial sensitivity. Genome sequencing was performed to validate phenotypic resistance profiles and determine the molecular characteristics of strains. CIA-R *E. coli* and *K. pneumoniae* were detected in 69 (30.1%) and 20 (8.73%) of the faecal swabs respectively. Two large urban locations tested positive for CIA-R *E. coli* (frequency ranging from 34.3%-84.3%), and/or for CIA-R *K. pneumoniae* (frequency ranging from 12.5%-50.0%). A small number of CIA-R *E. coli* (3/31, 9.7%) were identified at a small tourist town, but no CIA-R bacteria were recovered from gulls at remote sites. Commonly detected *E. coli* sequence types (STs) included ST131 (12.5%) and ST1193 (10.0%), and five *K. pneumoniae* STs were found. Resistance genes including *bla*_CTX-M-3_, *bla*_CTX-M-15_ and *bla*_CTX-M-27_ were identified in both bacterial species. High-level colonisation of CIA-R *E. coli* and *K. pneumoniae* in Silver Gulls in and around urban areas compared to remote locations substantiates that anthropogenic activities are strongly associated with acquisition of resistant bacteria by gulls.

**Importance:** Humans play an important role in dissemination of antimicrobial resistant bacteria. This study investigated the relationship between carriage rates of resistant bacterial pathogens *(Escherichia coli* and *Klebsiella pneumoniae*) among Silver Gulls and their proximity to human populations. The frequency of resistant *E. coli* carriage was high (ranging from 34.3 – 84.3%) in the samples collected from areas with high human population density while resistant *K. pneumoniae* frequencies at these sites varied from 0 to 50%. However, resistant *E. coli* and *K. pneumoniae* were not recovered from any of the remote sites that did not have a permanent human population. This study, conducted across a large stretch of the southwestern Australian coastline, indicated that seagulls act as vectors in carrying and disseminating antimicrobial resistant bacteria, including clinically significant strains. High-level colonisation of resistant *E. coli* and *K. pneumoniae* in Silver Gulls in and around urban areas compared to remote locations substantiates that human activities are strongly associated with acquisition of resistant bacteria by Silver gulls.

## Introduction

Globally, seagulls are thought to play a significant role in carriage and dissemination of pathogenic bacteria expressing resistance to critically important antimicrobials (CIA).^1,2,3^ In Australia, Silver Gulls (*Chroicocephalus novaehollandiae*) are common native fauna of coastal environments with a tendency to congregate as large flocks in urban areas owing to the successful adaptation of scavenger-based foraging. An Australia-wide survey of this species demonstrated that urban seagulls have high levels of faecal carriage of fluoroquinolone and extended-spectrum cephalosporin-resistant (CIA-R) *E. coli*, with prevalence rates of 24% and 22% respectively.^4^ The sequence types (STs) identified were human-associated extra-intestinal pathogenic *E. coli* ST131 (clades O25:H4 *H*30-R and *H*30-Rx) that is reported globally, ST1193, and other clinically significant strains belonging to ST10, ST69, ST38, ST95 and ST450. These are known, in humans, to cause severe life-threatening infections such as septicaemia, gastroenteritis, urinary tract infection, neonatal meningitis and hospital acquired pneumonia.^4^ The potential role of seagulls as a wildlife vector for the spread of antimicrobial resistance (AMR) was further substantiated by an investigation towards the transmission of resistant *E. coli* strains and associated mobile genetic elements (MGE) between different species of wild birds (including seagulls) sharing a coastal habitat adjacent to an urban environment.^5^ The study found a significantly higher frequency of CIA-R *E. coli* in seagulls (53%) compared to penguins (11%) and pigeons (10%) with no CIA-R *E. coli* found in Bridled Terns.

The widespread and elevated levels of carriage of CIA-R pathogenic *E. coli* of anthropogenic origins among seagulls in the absence of any direct exposure to antimicrobials raises concerns for public health since both the site of selection and source of exposure is uncertain.^6^ Sewage, landfill refuse facilities and effluent from hospitals and nursing homes are hypothesised to be involved as exposure sources owing to the seagull’s propensity to scavenge across varying urban environments. While proximity to human activities is considered to be the primary reason for the transference of CIA-R bacteria to seagulls ^7,8^, there is a lack of direct evidence in the form of comparison of rates of colonisation between birds found in areas of low and high density of human habitation.

One limitation of previous studies is that they focused solely on detection and evaluation of CIA-R *E. coli* carriage in wild birds and seagulls in particular. It is unclear if seagulls also carry other bacterial species expressing clinically important forms of resistance that also might act as a useful signal for ecological linkage between humans and seagulls. *Klebsiella pneumoniae* is another member of the Enterobacteriaceae family and is one of the ESKAPE pathogens (comprising *Enterococcus faecium, Staphylococcus aureus, K. pneumoniae, Acinetobacter baumannii, Pseudomonas aeruginosa* and *Enterobacter* species).^9^ *K. pneumoniae* is an opportunistic pathogen causing sepsis, urinary tract infections, cystitis, surgical wound infections, and septicaemia in humans, causing high mortality rates and extended hospitalization.^10^ Like *E. coli, K. pneumoniae* demonstrates a high propensity to acquire resistance to critically important antimicrobials, making it difficult to treat. Moreover, there is good evidence that resistance determinants readily transfer amongst the *Enterobacteriaceae* family, including *Klebsiella* and *E. coli*.^11^ *K. pneumoniae* is in fact regarded as adept at acquiring resistance to CIAs, including carbapenems and is viewed as a prominent carrier and disseminator of resistance genes to clinically significant human pathogens from various environmental sources.^12^

In this study we hypothesise that the carriage of CIA-resistance, via *E. coli* and *K. pneumoniae*, in gulls is related to the density of the local human population, and that the level of carriage decreases as distance from areas of human habitation increases. The validity of this hypothesis was tested by assessing a trend between the human population density and carriage of CIA-R *E. coli* and *K. pneumoniae* in seagulls.

## Materials and methods

### Sample collection

Samples were collected by swabbing freshly voided Silver Gull faecal droppings which then were transported in Ames Charcoal Media (Copan) to the Antimicrobial Resistance and Infectious Diseases Laboratory at Murdoch University for processing within four to five days of collection. A total of 229 samples were collected from 10 different southern coastline locations in Western Australia (WA). The sampling locations including main town centres and remote locations as shown in Table 1. Amongst the selected sampling locations, the Albany area (i.e. Albany township and adjacent Emu Point Beach) had the largest human population (36,583 as of 2016)^13^ and is WA’s sixth largest town. The Esperance area (i.e. Esperance town centre and nearby Bandy Creek) was identified as densely populated location, with a population size of 14,236 (2016 census).^13^ The township of Denmark has a relatively low base population size (5,845 as of 2016)^13^, although this increases by several times during the tourist season. All the other sites, comprising Conspicuous Cliff, Betty’s Beach, Cheynes Beach, Lucky Bay, and Cape Arid have no permanent residents, although people visit for tourism. They were classified as remote regions due to their lack of permanent residents and distance from the nearest town centres (>10 km). The sampling sites and population density are shown on the map of WA (Figure 1). This study was approved by Murdoch University Animal Ethics Office (Animal Ethics Cadaver/ Tissue Notification Permit No. 872).

**Table 1:** Sampling location details with number of swabs collected from each location

| Sampling Location | Region | Number of Swabs | Location Description | Distance from Main City/Townships | Human Population (2016) |
| --- | --- | --- | --- | --- | --- |
| Albany Town Centre | Urban | 32 | Port City | 418 km south east of Perth | 36,583 |
| Albany Emu Point Beach | Urban | 32 | Suburb and tourist destination | 8.5 km from Albany | 316 |
| Esperance Town Centre | Urban | 61 | Town | 720 km from Perth, 482 km from Albany | 14,236 |
| Esperance Bandy Creek | Urban | 10 | North eastern suburb of Esperance | 6 km from Esperance business district | 304 |
| Denmark | Semi-urban | 31 | Tourist township | 423 km south-south-east of Perth | 5,845 |
| Betty's Beach | Remote | 11 | Tourist destination | 50 km east of Albany | 0 |
| Cape Arid | Remote | 8 | Tourist destination | 731 km south east of Perth, 120 km east of Esperance | 0 |
| Cheynes Beach | Remote | 35 | Tourist destination | 65 km east of Albany and 470 km south of Perth | 0 |
| Conspicuous Cliff | Remote | 4 | Tourist destination | 13 km east of township of Walpole | 0 |
| Lucky Bay | Remote | 5 | Tourist destination | 63.6 km east of Esperance | 0 |

**Figure 1.**
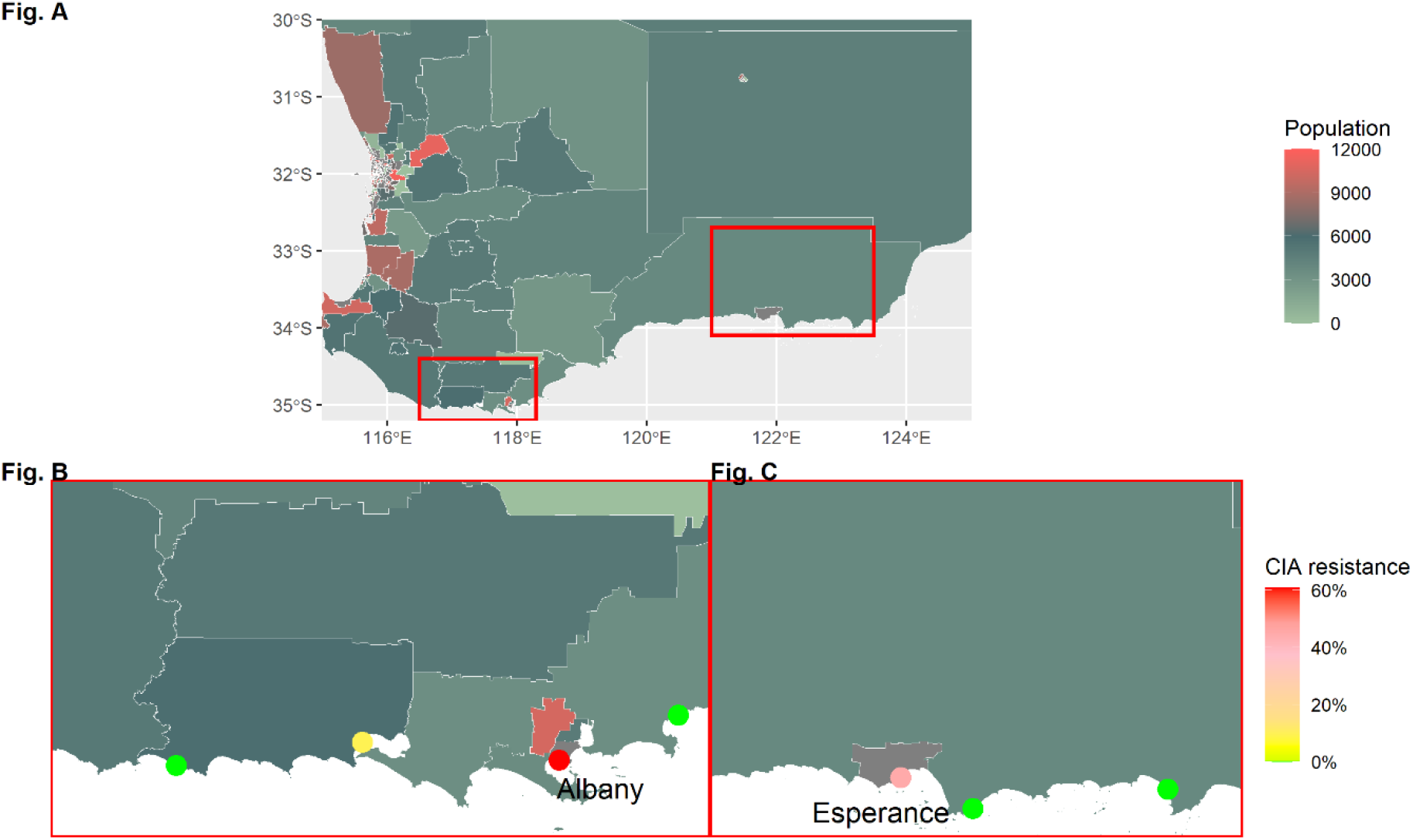
Proportion of CIA resistance found in seagull faecal droppings collected from different sampling location in Western Australia (WA). A is a choropleth map showing the population density for WA, with the red box showing the area where sampling was performed. The inset map “B” is zoomed in on the sampling areas, with the proportion of CIA resistance shown in points (green=0 to red=100%), while choropleths show the human population density. Resistance can be seen in the populated areas of Albany and Esperance.

### Isolation

The isolation procedure included initial enrichment of swabs in three mL of buffered peptone water (ThermoFisher) for four hours. The enriched samples were streaked onto two selective agar plates to isolate resistant *E. coli* and *K. pneumoniae*, and were incubated for 16-20 hrs at 37°C: These plates were respectively screened for presumptive ciprofloxacin-resistant (MacConkey agar infused with 1 ug/mL ciprofloxacin, ThermoFisher) and ceftriaxone-resistant (Brilliance ESBL, ThermoFisher) Enterobacteriacae colonies. Presumptive CIA-R *E. coli* and *K. pneumoniae* (one colony per species per plate) were further sub-cultured on Sheep Blood Agar plates (ThermoFisher) and species identity confirmed by matrix-assisted laser desorption ionization-time of flight mass spectrometry (MALDI-TOF MS) (Microflex, Bruker).

### Antimicrobial sensitivity testing

Minimum inhibitory concentrations (MIC) for antimicrobials were determined for the 29 *K. pneumoniae* and 98 CIA-R *E. coli* isolates recovered from the selective plates. Broth micro-dilution was performed using the Robotic Antimicrobial Susceptibility Platform (RASP)^14^ as per Clinical Laboratory Standards Institute (CLSI) guidelines, with recommended breakpoints for interpreting phenotypic antimicrobial resistance being applied.^15^ In addition to ciprofloxacin and ceftriaxone, the panel included antimicrobials of low and high importance for public health including ampicillin, gentamicin, sulfamethoxazole/ trimethoprim, and tetracycline. The control culture *E. coli* ATCC 25922 was used as per CLSI guidelines.^15^ The MIC data were analysed using the EUCAST epidemiological cut-off value (ECOFF) wildtype breakpoints as indicators of resistance.

Samples were confirmed as positive for CIA-R *E. coli* or *Klebsiella* if they yielded growth of an *E. coli* or *Klebsiella* on the screening plates (based on species confirmation by MALDI-TOF MS) and demonstrated phenotypic resistance based on ECOFF to a either ciprofloxacin or ceftriaxone in the broth microdilution assays.

### Whole genome sequencing

Whole genome sequencing (WGS) was performed on a subset of confirmed CIA-R *K. pneumoniae* (n=14) and CIA-R *E. coli* (n=40) isolates based on susceptibility profiles. DNA extraction was performed from isolate cultures grown overnight on Sheep Blood Agar by using the MagMax DNA Multi-Sample extraction kit (ThermoFisher Scientific) according to manufacturer’s protocol. Sequencing libraries were prepared using the Celero DNA-seq library preparation kit (NuGEN) according to the manufacturer’s protocol. Sequencing was performed on the Illumina NextSeq 550 platform using a Mid-Output 300 cycles Kit v2.5.

*De novo* assembly of the sequence data was performed using SPAdes v3.14.0.^18^ Resistance and virulence genes were identified using ABRicate v1.0.1 ^19^ with ResFinder ^20^ and VFDB ^21^, based on the *de novo* assembled draft genomes. Identified resistance and virulence genes were considered present if they were at greater than 95% coverage and identity. The structure of contigs harbouring resistance genes identified in both species were investigated further in Geneious Prime v2021. Multi-locus sequence types (ST) for each isolated were identified using the MLST tool (version 2.19.0) described by Torsten Seeman using pubMLST data base (Seemann, 2019).^22^

## Results

### Detection of CIA-R *E. coli* and *K. pneumoniae*

A total of 229 swabs were collected from selected site. We found a total of 98 *E. coli* isolates and 27 *K. pneumoniae* isolates were recovered from the selective agar plates, but not all were confirmed to be CIA-R (i.e. resistant to ciprofloxacin and/or ceftriaxone) following microbroth MIC testing. Among the 229 swabs collected, 30.1% (n=69) were confirmed positive for CIA-R *E. coli* and 8.73 % (n=20) for CIA-R *K. pneumoniae*. Only 7.42% (n=17) of the swabs yielded both CIA-R *E. coli* and *K. pneumoniae*. None of the samples from remote areas yielded CIA-R *E. coli*. Seagull samples from the urban locations were positive for CIA-R *E. coli* at frequencies ranging from 34.3 -84.3% (Table 2). A relatively low rate of CIA-R *E. coli* (3/31; 9.7%) was detected in the small semi-urban tourist town of Denmark (Table 2). CIA-R *K. pneumoniae* was detected only in the urban areas of Esperance and Albany, with frequencies ranging from 12.5%-50.0% (Table 2). The overall proportion of resistance against CIAs was clearly higher in regions with higher human population density (Figure 1 and 2).

**Table 2:** Percentage of *E. coli* and *K. pneumoniae* isolated from seagull faecal droppings collected from different sampling location in Western Australia. The table shows total number of swabs collected from targeted regions and total confirmed CIA resistance in both *E. coli* and *K. pneumoniae* against ciprofloxacin (CIP), ceftriaxone (ESBL), and ciprofloxacin and/or ceftriaxone (CIA)

| Location | Region | Swabs (n) | Number (n) of swabs positive (%) |  |  |  |  |  |
| --- | --- | --- | --- | --- | --- | --- | --- | --- |
|  |  |  | *E. coli*_CIP (n=64) | *E. coli*_ESBL (n=44) | *E. coli*_CIA (n=69) | *K. pneumoniae*_CIP (n=17) | *K. pneumoniae*_ESBL (n=19) | *K. pneumoniae*_CIA (n=20) |
| Albany Town Centre | Urban | 32 | 26 (81.2) | 22 (68.7) | 27 (84.3) | 2 (6.2) | 4 (12.5) | 4 (12.5) |
| Albany Emu Point Beach | Urban | 32 | 11 (34.3) | 2 (6.2) | 11 (34.3) | 0 | 0 | 0 (0) |
| Esperance Town Centre | Urban | 61 | 18 (19.6) | 16 (22.9) | 22 (36.1) | 10 (11.4) | 10 (11.4) | 11 (18.0) |
| Esperance Bandy Creek | Urban | 10 | 6 (60) | 4 (40) | 6 (60) | 5 (50) | 5 (50) | 5 (50) |
| Denmark | Semi-Urban | 31 | 3 (9.7) | 0 | 3 (9.7) | 0 | 0 | 0 |
| Betty's Beach | Remote | 11 | 0 | 0 | 0 | 0 | 0 | 0 |
| Cape Arid | Remote | 8 | 0 | 0 | 0 | 0 | 0 | 0 |
| Cheyne's Beach | Remote | 35 | 0 | 0 | 0 | 0 | 0 | 0 |
| Conspicuous Cliff | Remote | 4 | 0 | 0 | 0 | 0 | 0 | 0 |
| Lucky Bay | Remote | 5 | 0 | 0 | 0 | 0 | 0 | 0 |

**Figure 2.**
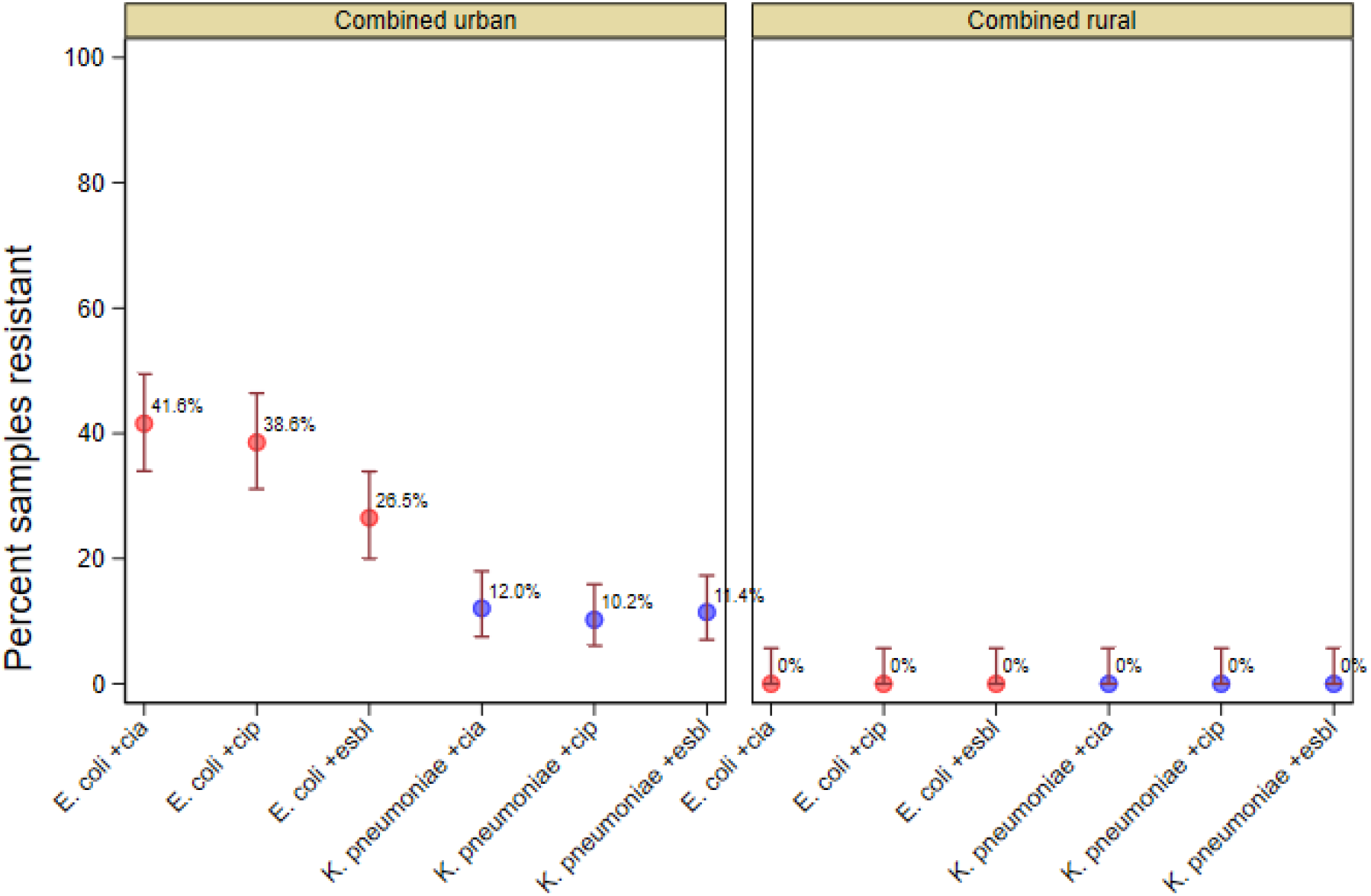
Percent of seagull faecal swab positive (± 95% confidence intervals) for *E. coli* (red marker) and *K. pneumoniae* (blue markers) expressing resistance to ciprofloxacin (+CIA), extended spectrum beta lactams (+ESBL) and critically important antimicrobials (+CIA, either +CIP or + ESBL) with data combined for all urban (including semi-urban) and all remote locations.

### Phenotypic and genotypic characteristics

#### *E. coli* isolates

Minimum inhibitory concentration (MIC) testing against a panel of six antimicrobials was performed on 98 *E. coli* isolates recovered from the selective plates. Resistance to ceftriaxone and ciprofloxacin was confirmed in 51% and 84.7% of the isolates respectively. High levels of resistance to ampicillin (98.0%), tetracycline (64.3%) and sulfamethoxazole/trimethoprim (51%) were found, with 17.3 % of isolates demonstrating resistance towards gentamicin (Figure 3).

**Figure 3:**
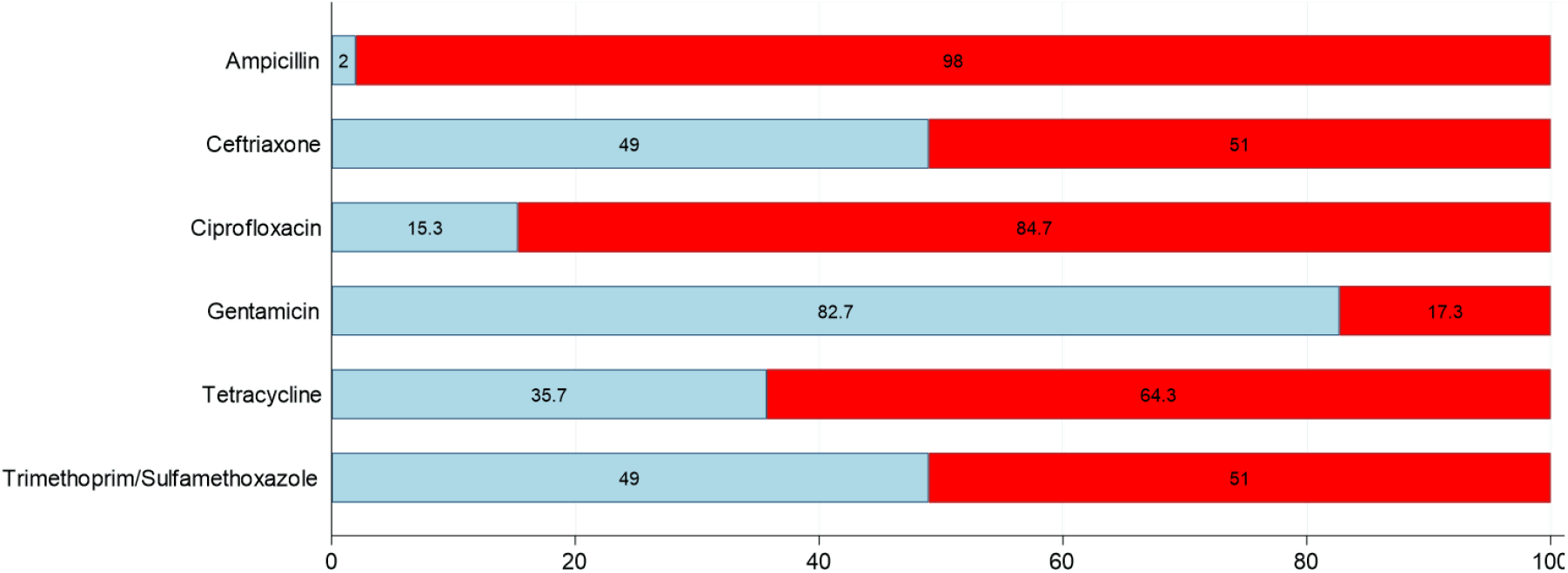
Overall resistance of *E. coli* isolates recovered from seagulls on CIA-resistance screening agar plates from two urban and a semi-urban location in Western Australia. Interpretation is based on EUCAST ECOFF wildtype breakpoints. No CIA-R isolates were found in remote locations. A single isolate per sample was included in the analysis. Key: blue – percent wildtype, red - percent non-wildtype (resistant).

Whole genome sequencing on a representative subset of CIA-R *E. coli* (n=40) isolates revealed 28 different sequence types (STs) (Table 3). The most frequently detected were ST131 (12.5%) and ST1193 (10.0%) (Table 3). Beta-lactam resistance genes were found in all but one of the sequenced *E. coli* isolates, with *bla*_TEM-1_ being the most commonly found (50.0%), followed by *bla*_EC-5_ (27.5%), *bla*_CTX-M-15_ (22.5%) and *bla*_CTX-M-27_ (12.5%). Tetracycline resistance-associated genes were found among 64.1% of the isolates, with *tet*(A) found most frequently (27.5%) followed by *tet*(B) (25.0%), and *tet*(D) and *tet*(M) (both at 5%). A single isolate carried *bla*_CMY-42_. Plasmid Mediated-Quinolone Resistance (PMQR) genes *qnr*S1 and *qnr*B4 were detected in 22.5% and 7.5% of the *E. coli* isolates, respectively. However, only two isolates with PMQR were associated with phenotypic resistance against ciprofloxacin.

**Table 3.**
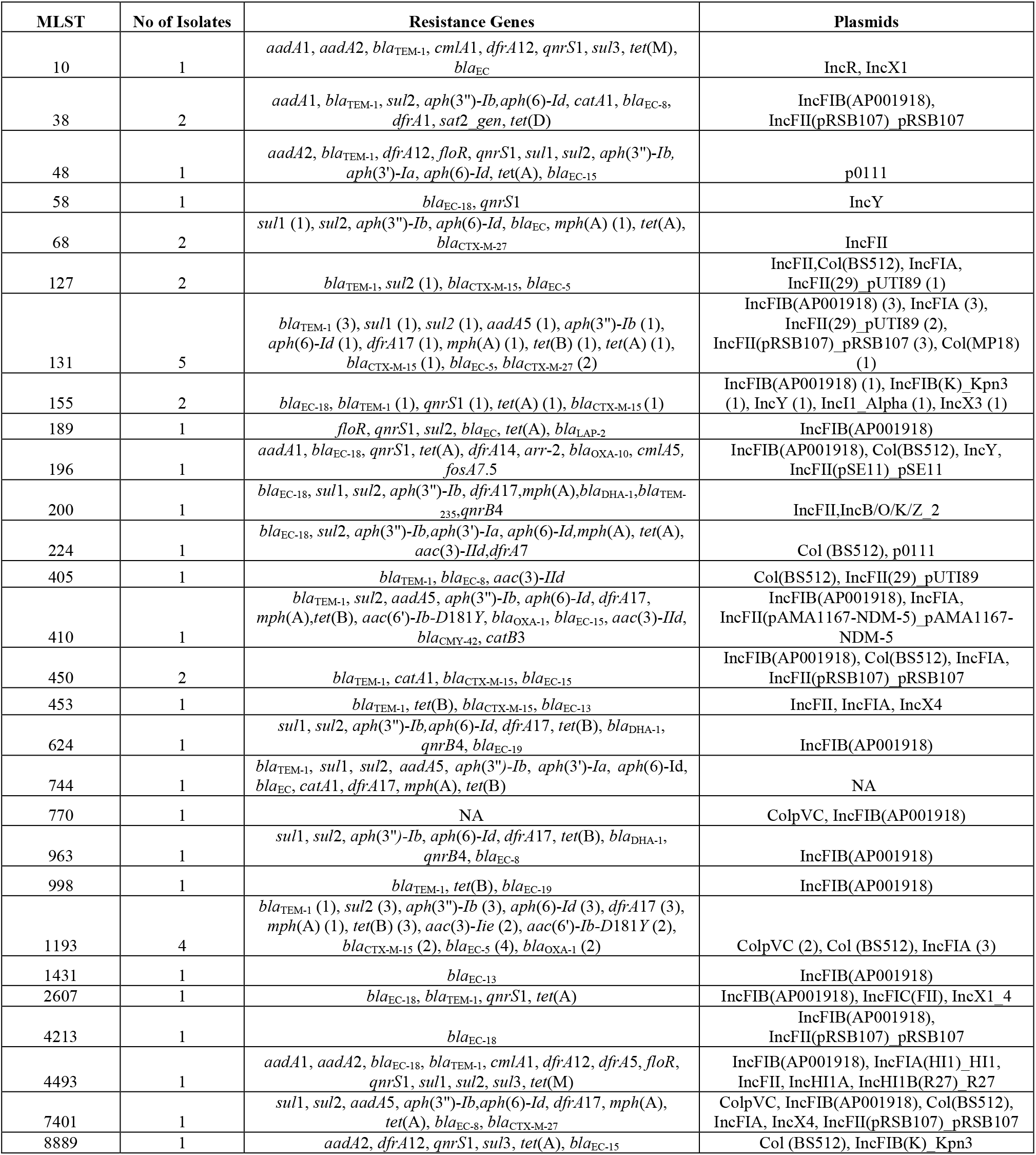
Number of *E. coli* isolates identified according to sequence type in MLST, with their associated resistance genes and plasmids

Using PlasmidFinder, a total of 24 different plasmids were predicted among 39 of the 40 *E. coli* isolates. The most commonly found plasmids were IncFIB(AP001918) (50.0%), Col(BS512) (32.5%), IncFIA (32.5%) IncFII(pRSB107)_pRSB107 (22.5%) and IncFII (17.5%). ST131 carried the highest number of different plasmids (30.0%) when compared to other STs such as ST1193 (22.5%), ST450 (20.0%), and ST127 (17.5%).

#### *K. pneumoniae* isolates

*K. pneumoniae* isolates (n=27) exhibited high frequencies of resistance to ampicillin (100%), sulfamethoxazole/trimethoprim (100%), tetracycline (81.5%), ceftriaxone (88.9%) and ciprofloxacin (85.2%) (Figure 4). Only 18.5% of *K. pneumoniae* isolates were resistant to gentamicin.

**Figure 4:**
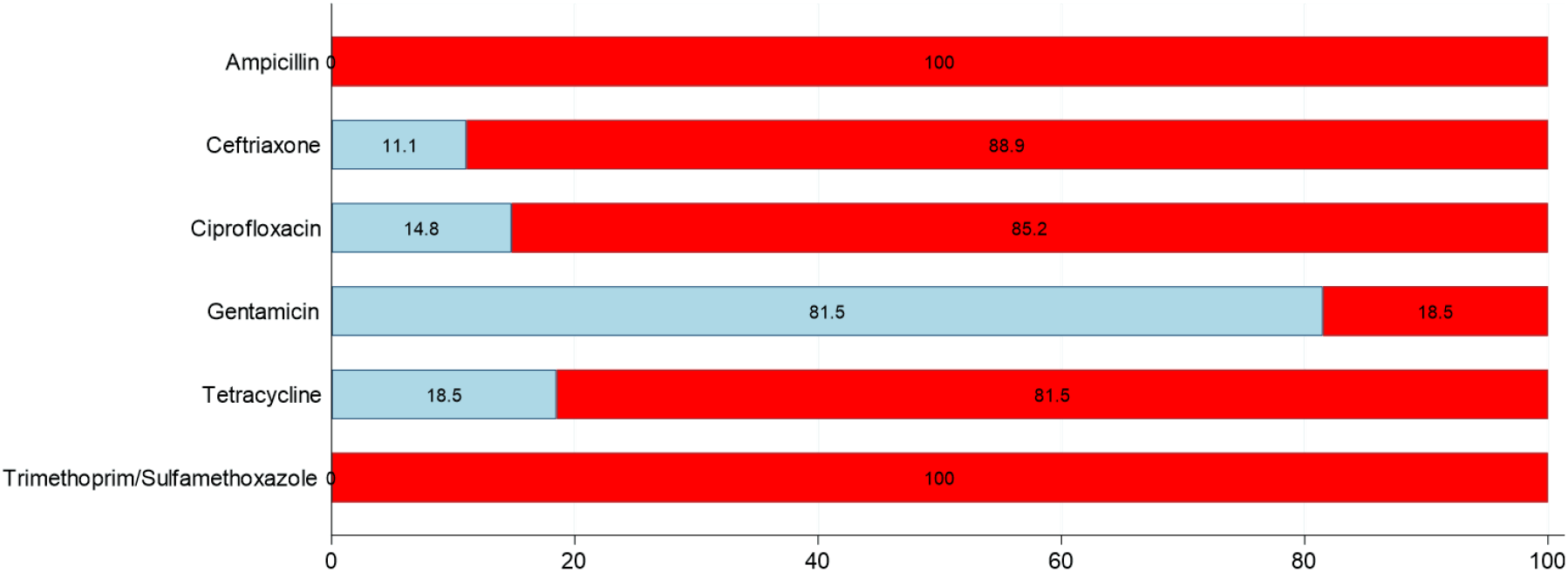
Overall resistance of *K. pneumoniae* isolates from seagulls on CIA-resistance screening agar plates from urban locations in Western Australia. Interpretation is based on EUCAST ECOFF wildtype breakpoints. Non-wildtype isolates were only found at Albany town centre and at Esperance town centre and nearby Bandy creek. A single isolate per bird was included in the analysis. Key: blue – percent wildtype, red - percent non-wildtype (resistant).

Genome sequencing of a subset of 14 CIA-R *K. pneumoniae* isolates identified 5 different ST types, comprising ST4568 (64.3%), ST6 (14.3%) and ST485, ST967 and ST307 (all at 7.1%). Of the beta-lactamase genes, *bla*_TEM-1_ was the most frequently detected (92.8%), followed by *bla*_CTX-M-3_ (71.4%), *bla*_SHV-187_ (71.4%) and *bla*_CTX-M-15_ (21.4%). Fosfomycin resistance genes (*fosA*) were also found in all isolates. Tetracycline resistance genes including *tet*(A), *tet*(B), *tet*(D), *tet*(M) were found in 78.5% of the selected isolates. Macrolide resistance genes (*mph*) were identified among 71.4% of isolates. PMQR genes were detected in 13 isolates, including *qnr*S1 (78.5%) and *qnr*B1 (14.2%). Only two isolates carried two different plasmids (ColRNAI, IncFIB(K)_Kpn3).

## Discussion

The aim of the study was to investigate whether there was a relationship between the carriage of CIA-R bacteria by gulls and the density of the human population in the sampling areas. There are a number of critically important antimicrobials,^23^ but this study focused on resistance to two of the most important classes – quinolones (exemplified by ciprofloxacin) and 3^rd^ and 4^th^ generation cephalosporins (exemplified by ceftriaxone). The findings indicate a strong association between nearby human population and carriage of CIA-R bacteria by gulls. The frequency of CIA-R *E. coli* carriage was high in the samples collected from the populated areas of Albany Town Centre and nearby Emu Point, and in Esperance Town Centre and nearby Bandy Creek (ranging from 34.3 – 84.3%). *K. pneumoniae* CIA-R frequencies at these sites varied from 0 to 50% (Figure 1 and 2). The short distances between the town centres and nearby beaches mean that local gull population could easily move between them.^24^ In contrast, at the small semi-urban tourist township of Denmark the carriage rate of CIA-R *E. coli* was much lower (9.7%), and no CIA-R *K. pneumoniae* was identified. Most importantly, CIA-R bacteria were not recovered from any of the remote sites that did not have a permanent human population (Figure 1 and 2). Accordingly, there was a clear correlation between the presence and density of human population and the occurrence of CIA-R *E. coli* and *K. pneumoniae* carriage by local gulls (Figure 1 and 2). Fewer samples were obtained at the remote sites compared to the urban sites, and this reflected a much lower density of gull populations at the remote sites – meaning that it was more difficult to obtain larger numbers of fresh representative samples. The higher populations of gulls at the urban sites is likely to reflect more abundant food sources, but also increases the likelihood of transmission of CIA-R bacteria between the birds and into the local environment where humans may be exposed. It is somewhat reassuring that the gulls in the remote sites away from human populations did not carry detectable CIA-R bacteria; however, they were only screened for *Enterobacteriaceae* possessing two classes of CIA, and it is possible that strains carrying resistance to other CIAs went undetected.

Apart from ciprofloxacin and ceftriaxone, the other antimicrobials selected for MIC testing included ones that are also categorised as critically and highly important by World Health Organisation.^23^ These antimicrobials belong to the major drug classes and included Ampicillin (aminopenicillins), Gentamicin (aminoglycosides), Tetracycline (chlortetracycline) and sulfamethoxazole/trimethoprim (sulfonamides, dihydrofolate reductase inhibitors and combination).^23^ Rates of resistance to these antimicrobials were also high in the isolates recovered from urban areas, but their occurrence in remote areas could not be tested because no non-CIA-R isolates were available as a result of the screening method used.

Although the findings provide new details on the ecological distribution of CIA-R human pathogens in urban wildlife, there remains uncertainty about the consequence of this for in-contact human populations or for colonisation of food-producing animals by exposure to gulls scavenging around livestock production enterprises. Seagulls are frequent visitors to the food-animal enterprises in some remote areas, and so in these circumstances’ transmission of CIA-R *E. coli* or *K. pneumoniae* to animals via contamination of pasture, food troughs and/or watering points is a real prospect. In both cases there is scope for newly acquired CIA resistance from the environment to be amplified by use of antimicrobials for treatment of human or food-animal disease. There also is potential for detection of such organisms during surveillance or diagnostic investigation to be inappropriately attributed to emergence of CIA resistance in the corresponding host due to poor antimicrobial stewardship.

While the current study evaluates a large section of coastline comprising approximately 650 km of seagull habitat, this is only small in relation to the entire coastline of Australia, and so there is a need to establish if the relationship between humans and CIA-R carriage in seagulls is more generally applicable. A barrier to working across such a vast expanse has hitherto been the capacity of laboratories to process meaningful numbers of samples and assessing sufficient isolates in adequate details. With the evolution of RASP technology, demonstrated in this study, it is clear that high throughput processing of colonies by laboratory robotics can succeed without sacrifice of measurement quality, thus making it possible now to undertake a continental-scale comparison of the CIA-R bacteria carriage rates in urban seagulls and humans.

This study further substantiates that avian species like gulls are not only limited to accumulating, amplifying, and disseminating CIA-resistant *E. coli*, but they can also disseminate other key Gram-negative bacteria that are pathogenic for humans, such as *K. pneumoniae* which is known for being highly virulent with a propensity to cause invasive infections.^25^ The latter organism is highly adept at rapidly accumulating multi-drug resistance genes including those responsible for producing *K. pneumoniae* carbapenemase (KPC), New Delhi metallo-β-lactamase-1 (NDM-1) and other β-lactamase enzymes that make treatment of infections highly challenging due to lack of alternative options.^11^ Out of the five different *K. pneumoniae* STs identified in this study, ST485 and ST307 previously have been detected in wastewater treatment plants (WWTP) and clinical samples in other parts of the world.^26^ *K. pneumoniae* ST307 has been described as a globally emerging lineage carrying multiple resistance plasmids transmissible to other *K. pneumoniae* STs, as well as to different bacterial species.^27, 28^ ST307 isolates in this study harboured the clinically significant human-associated resistance gene *bla*_CTX-M-15_. *K. pneumoniae* ST4568 was identified in this study, and although there is limited data on this organism’s impact on humans, it was found to possess resistance genes such as *bla*_CTX-M-15_ and *bla*_CTX-M-3_.

The current study recorded similar observations as in a previous Australia-wide investigation of the carriage of CIA-R *E. coli* amongst seagulls. These included finding a high prevalence of *E. coli* ST131, a virulent human strain associated with fluoroquinolone FQ resistance and *bla*_CTX-M-15_ type ESBL production, followed by rapidly emerging extraintestinal pathogenic *E. coli* (ExPEC) ST1193 and the presence of human associated resistance genes in these isolates.^4^ There are several other studies supporting the premise that anthropogenic contributions into the environment greatly influence the prevalence and carriage of antibiotic resistant bacteria by seagulls that inhabit the environment,^29-34^ potentiated by the foraging habits of seagulls.

## Conclusions

This study, conducted across a large stretch of the southwestern Australian coastline, has corroborated an earlier, smaller scale study that indicated that seagulls act as very efficient vectors in carrying and disseminating antimicrobial resistant bacteria, including clinically significant strains. Where the potential for interactions between humans and seagulls is greatest, colonisation with hazardous organisms occurs more frequently. Considering the ecological mobility of Silver Gulls and similar avian species, the findings point to a need to obtain a broader understanding of the pathways through which the resistant bacteria or genes enter these hosts and their subsequent fate in the environment following shedding in faecal material. Evidence is needed to understand and evaluate the need for measures that disrupt the pathway of contamination by reducing the accessibility of these species to ‘exposure hot spots’ such as human waste, hospital effluent and landfill sites.

## Funding

This project was supported by University of Adelaide and Murdoch University

## Transparency declarations

None to declare

